# Optimized Multiple Circular Sequence Alignment for Cyclic Peptide Motif Discovery

**DOI:** 10.64898/2026.07.28.741376

**Authors:** Ye Yuan, Zhe Li, Kaiqiang Hu, Pengwei Pan, Fang He

## Abstract

Head-to-tail (H2T) cyclized peptides are an increasingly important modality in drug discovery, combining high target affinity and selectivity with metabolic stability. Because their underlying chemistry is still that of a linear amino-acid chain, their linear sequence representation is the native input format of mainstream sequence generative models now driving de novo peptide design (Slough et al., 2018; Rettie et al., 2025a;b). Discovering the conserved motifs responsible for a family’s function across a *library* of such candidates requires a multiple sequence alignment (MSA). Because a cyclic peptide can be linearised at any residue, the alignment must additionally solve for the unknown rotation of each sequence, which is the multiple circular sequence alignment (MCSA) problem. However, leading MCSA heuristics (e.g. Ayad & Pissis, 2017) were tuned for the genomic regime (a few tens of long sequences) and become prohibitively slow on the cyclic peptide library regime (hundreds to thousands or more shorter sequences). We close this gap by identifying quality-preserving optimisation opportunities, notably the *library-scale preset* tailored to short-sequence inputs (algorithmic details in the Supplementary Material), and by adding an orthogonal multi-core backend for further performance tuning. We validate the pipeline on a library of 1,000 H2T cyclized peptides of length 18 targeting the oncoprotein Mouse double minute 2 human homolog (MDM2) produced by an internal peptide-design engine. In this practical setup, the optimised MCSA recovers the underlying positional motif of MDM2 binders at the same fidelity as the original MCSA implementation while running over 650× faster. Our optimised MCSA tool thus enables library-scale cyclic peptide sequence alignment and is publicly available at https://github.com/IVB-Generative-Biology/mars-turbo.

## 1 Introduction

Head-to-tail (H2T) cyclized peptides, macrocycles whose N- and C-termini are joined by an amide bond, are a prominent modality in modern drug discovery. Cyclisation rigidifies the backbone, improving target affinity and selectivity and protecting against proteolytic degradation, while the molecule’s underlying chemistry remains that of a linear amino-acid chain (Slough et al., 2018). This property makes H2T peptides especially attractive computationally: their linear sequence representation is the native input format of the mainstream sequence generative models now driving de novo peptide design, from AlphaFold2-based designers such as AfCycDesign to RFdiffusion-derived macrocycle generators such as RFpeptides (Rettie et al., 2025a;b).

A frequent downstream task on a *library* of such computationally- or experimentally-generated candidates is motif discovery: locating the conserved residues and positions that recur across the family and are therefore likely responsible for its function. The standard route to a motif is a multiple sequence alignment (MSA), from which a consensus column and a position-wise conservation profile are read off (D’haeseleer, 2006). Throughout we write

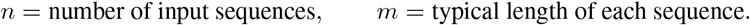

For the genomic applications that originally motivated alignment tooling, *n* is small (a few tens of genomes) and *m* is large (tens of thousands of bases). For a cyclic peptide drug-discovery programme the situation is reversed: *n* runs to hundreds to thousands or more candidates per library, while *m* is the short, fixed length of the designed macrocycle. Tools that scale well in *n* are therefore essential.

MSA alone is insufficient for cyclic peptides, because a cyclic molecule can be linearised at any of its *m* residues and the choice is a modelling convention rather than a property of the molecule. Standard MSA assumes the left- and right-most positions are meaningful, so applying it directly to arbitrarily-linearised cyclic sequences misaligns homologous positions. The same issue affects mitochondrial, viroid, and viral genomes linearised at arbitrary points in population studies, and is reflected in inconsistent linearisation standards across sequence databases (Ayad & Pissis, 2017). *Multiple circular sequence alignment* (MCSA) restores correctness by additionally choosing, for each input sequence, a rotation (cyclic shift) that yields a lower-distance alignment.

MARS (Ayad & Pissis, 2017) is the leading MCSA heuristic. It runs in three stages: **(1)** a pairwise comparison estimates, for every pair of sequences, an approximate cyclic distance and a best relative rotation, which is then refined by a banded alignment of the sequence ends (Needleman & Wunsch, 1970); **(2)** a guide tree is built from these distances (Saitou & Nei, 1987); **(3)** profiles are aligned progressively down the tree, re-rotating at every merge (Feng & Doolittle, 1987; Thompson et al., 1994). The first two stages scale quadratically and cubically with the number of sequences, respectively. Both are well matched to the genomic regime (a small number of sequences) but become the bottleneck for a peptide library (a large number of sequences): in our experiments the original implementation already requires several minutes on only 500 sequences of length 2,000.

### Contributions

We engineer MARS to scale to peptide libraries while preserving output quality, and validate the result rigorously.

- **A reference-based rotation-recovery scheme specific to circular sequences**. The rotations of all pairs are not independent: each sequence has a single absolute frame, so a sequence’s rotation relative to any other follows by transitivity once its frame is fixed. We therefore recover each frame from only a handful of reference comparisons instead of comparing all pairs. To our knowledge this structure has not been exploited before, and it is what lets the pairwise stage run in time near-linear in the number of sequences rather than quadratic (Supplementary Material, Fig. S2).
- **Three quality-preserving optimisations** that remove the stages scaling poorly with the number of sequences: skipping a redundant refinement step, estimating pairwise relations from a few references (above), and using a faster guide tree together with a reorganised scoring routine that is algebraically exact. Each preserves output quality; full pseudocode is in the Supplementary Material (Figs. S2–S5).
- A controlled evaluation on the original benchmark suite (Ayad & Pissis, 2017) (synthetic and real datasets), using the published alignments as an exact ground-truth anchor for the average-pairwise-distance metric, with every timing measured across three repetitions.
- An orthogonal multi-core and SIMD (single instruction, multiple data) backend that is bit-identical to the single-thread run and scales near-linearly with core count on the pairwise-comparison stage.

Together these yield one to two orders of magnitude of end-to-end speedup, seen as the expected scaling improvement (quadratic/cubic to near-linear in the sequence count) rather than a machine-dependent constant factor, with output quality preserved by the library-scale preset on all but the shortest, most divergent sets. We frame the method for cyclic peptide motif discovery and demonstrate it on the original genomic benchmark, on scaled synthetic sets that isolate the large-library regime, and on a dedicated H2T cyclic peptide library (§3.3).

## 2 Materials and Methods

### 2.1 Experimental and technical design

The overall design of the study is summarised in the Supplementary Material (Fig. S1). We take the original MARS implementation as an exact reference and evaluate three families of inputs: the twelve datasets of the original benchmark, which anchor our tool to the published results; scaled synthetic datasets (100–500 sequences of length 2,000) that isolate the large-library regime; and a dedicated H2T cyclic peptide library (1,000 length-18 sequences targeting MDM2) produced by an internal design engine. Every claim is made against this reference through two validation gates, calibration and reproduction (§2.4), with alignment quality read out through the AVPD metric and circular frame consistency (§2.2), and with every timing averaged over three repetitions. The optimisations (§2.3) are grouped into command-line presets (§2.5), so that the quality–speed trade-off is a single user choice.

### 2.2 AN ACCURACY CRITERION: CIRCULAR FRAME CONSISTENCY

MARS’s output is a set of rotated sequences, so the natural correctness question is *not* “do the rotations match a reference bit-for-bit”. Two factors make exact rotation matching a noisy yardstick. First, a global cyclic shift applied to *every* sequence is an equivalent circular alignment, so there is no privileged start. Second, NJ guide trees contain many near-ties, so the specific frame a run settles on is unstable even at unchanged quality. The correct criterion is **circular frame consistency**: do the chosen rotations place homologous fractional positions into a common frame? We measure this (for our controlled synthetic data, where a ground-truth frame is known) as the circular variance of the per-sequence rotation phases; lower is better, with random rotations scoring *≈*0.95 and a consistent frame scoring *≈*0. This mirrors the well-established finding that MSA *quality* is robust to the specific guide tree even when the alignment differs (Blackshields et al., 2006; Edgar, 2010).

### 2.3 OPTIMISATIONS

We target the three stages of MARS that scale poorly with the number of sequences: the pairwise comparison, the pairwise refinement, and the guide tree, together with the per-column cost of the progressive alignment. Each change preserves output quality; the precise mechanisms and pseudocode are deferred to the Supplementary Material (Figs. S2– S5), and the command-line presets that select combinations of them are described in §2.5.

#### A. Skip a redundant refinement step

Stage 1 refines every pairwise rotation, but Stage 3’s progressive alignment re-refines rotations at every merge anyway. The first pass is therefore redundant; this is the same observation that lets Clustal and MAFFT omit it (Thompson et al., 1994; Katoh & Standley, 2013), and we drop it.

#### B. Estimate pairwise relations from a few references

Instead of comparing every pair of sequences, we compare each sequence against only a handful of *reference* sequences. The pairwise *distances* for the guide tree come from a landmark embedding, the scheme behind Clustal Omega’s fast guide tree (Blackshields et al., 2010), which is adequate because alignments are robust to the guide tree (§2.2). The *rotations* exploit a structure specific to circular sequences: each sequence has a single absolute frame, so once a sequence’s frame is recovered from the references, its rotation relative to any other sequence follows by transitivity. On our data the recovered rotations match the true ones within a few bases. This is the key idea that lets the pairwise stage run near-linearly rather than quadratically in the sequence count.

#### C. A faster guide tree and a reorganised scoring routine

We replace the cubic-time guide tree with a quadratic-time one (Sokal & Michener, 1958), again acceptable because the tree is quality-tolerant (§2.2). We also reorganise the progressive alignment’s per-column scoring so that repeated work is done once; this reorganisation is algebraically exact, so the alignment output is bit-identical to the original.

#### D. Multi-core and SIMD acceleration.

The changes above reduce the total work; a second, orthogonal axis reduces wall-clock time through multi-core parallelism (OpenMP, OpenMP Architecture Review Board, 2018) and vectorised instructions (SIMD). Every independent stage is parallelised, and the parallel output is bit-identical to the single-thread run on every dataset. A portable scalar build is retained as a fallback (§2.5).

Taken together, every stage that was quadratic or cubic in the sequence count becomes near-linear or quadratic, while the progressive alignment (which is already linear in the sequence count) becomes the dominant cost. This is why the empirical speedups of §3.2 plateau once they reach that stage (Figure 2). Precise per-phase complexities are given in the Supplementary Material (Table S1).

### 2.4 Evaluation setup

#### Datasets.

We use the *exact* datasets the original authors used to benchmark MARS (Ayad & Pissis, 2017), cloned from their repository: nine synthetic INDELible (Fletcher & Yang, 2009) / Jukes–Cantor sets {12, 25, 50} ×2500 × {5, 20, 35} % (notation `n.length.div`), fed to MARS as the randomly rotated inputs, plus three real sets: Mammals (12 mtDNA, *∽*16.8 kbp), Primates (16 mtDNA, *∽*16.6 kbp), and VIROIDS (18 RNA, *∽*363 bp). To expose the regime the optimisations target (the cyclic peptide library scale, where the sequence count runs to hundreds rather than dozens), we add scaled synthetic sets with *n ∈ {*100, 250, 500} sequences of length 2000, at moderate (*∽*8%) and high (*∽*16%) divergence, simulated along random binary trees with random cyclic linearisation and a known ground-truth frame. MARS parameters follow the paper: synthetic/scaled *q*=5, *l*=50, *P* =1.0; real mtDNA *q*=5, *l*=100, *P* =2.0; Viroids *q*=4, *l*=25, *P* =1.0. We additionally evaluate on a dedicated H2T cyclic peptide library (1,000 peptides of length 18 targeting MDM2, produced by an internal peptide-design engine, each cyclically rotated by an independent random offset to pose a genuine frame-recovery task; *q*=3, *l*=6, *P* =1), reported in §3.3. The present evaluation establishes the algorithmic claims on the original genomic benchmark and on the scaled synthetic sets that isolate the relevant large-library regime.

#### Quality metric: AVPD

Our headline metric is the average pairwise distance (AVPD): the mean, over all unordered sequence pairs, of the number of alignment columns at which the two sequences differ (gaps counted as differences). This is the metric the original paper reports (its “average pairwise distance”), computed on the final MSA from ClustalΩ (Sievers et al., 2011) (v1.2.4). Lower is better. For scaled sets we additionally report *circvar* (circular frame consistency vs the known ground-truth frame; 0 = perfect) and the *rotation agreement* (*rotagr*), the fraction of sequences whose recovered rotation matches the reference build up to the single arbitrary global frame (1.0 = identical).

#### Two validation gates

To make the comparison credible we verify two things *before* reporting results. **Calibration:** our AVPD formula reproduces the original paper’s Table 5 *exactly* on the published MUSCLE alignments: Mammals 5421.04 (paper 5421), Primates 4070.98 (4070), Viroids 221.34 (221). **Reproduction:** our reference build reproduces the published `.mars.fas` refined outputs exactly (12/12, 50/50, 12/12; the few non-exact cases are circularly-identical rotation ties). Our build is therefore equivalent to the original MARS and serves as a valid reference.

#### Timing protocol

Every MARS run is repeated three times and we report the mean; run-to-run spread was small throughout (worst case under 9% on every timed cell, under 2% for the single-thread baseline). ClustalΩ is deterministic and is run once per cell with a 30-minute cap, and all cells completed. The multi-core (§3.4) and cyclic peptide (§3.3) experiments follow the same protocol. Full per-cell timing statistics are given in the Supplementary Material.^1^

### 2.5 Practical usage and presets

The algorithmic changes of §2.3 are exposed through a small set of command-line options, collected here so they do not clutter the algorithmic description. They are opt-in (the default build reproduces the original MARS); the multi-core/SIMD backend of optimisation D is always-on.

#### Quality presets (**-Q**)

A single flag selects combinations of the optimisations: `-Q 0` is the original MARS (no change); `-Q 1` applies (A) + (C2) (it drops the redundant refinement and uses the UPGMA tree) and is the recommended default, referred to throughout as the *library-scale preset*; `-Q 2` additionally enables the landmark q-gram (B) with *R*=5 references; `-Q 3` is the most aggressive, setting *R*=1. Each optimisation can also be toggled individually: `--no-refine` (A),`--qgram-refs R` (B), and `--guide-tree 1` (C2).

#### Threads and the portable build

`-T p` sets the number of OpenMP threads (optimisation D). A portable scalar build, obtained with `make NOAVX=1`, disables the SIMD profile-profile kernel and is retained as a fallback on architectures without SIMD support; it is otherwise bit-identical to the default build.

## 3 Results

### 3.1 Benchmark on the original MARS DATASETS

Tables 1 and 2 report quality (AVPD) and wall-time on the 12 datasets of the original benchmark.

**Table 1.** Quality (AVPD) on the original benchmark datasets. noMARS = ClustalΩ on unrefined input; Δ% = change vs the reference (-Q 0; negative = better than original MARS).

| Dataset | noMARS | $-Q\ 0$ | $-Q\ 1$ | | $-Q\ 2$ | | $-Q\ 3$ | |
| --- | --- | --- | --- | --- | --- | --- | --- | --- |
| | | | AVPD | $\Delta\%$ | AVPD | $\Delta\%$ | AVPD | $\Delta\%$ |
| 12.2500.5 | 1244.4 | 197.6 | 192.1 | -2.8 | 191.2 | -3.2 | 191.2 | -3.2 |
| 12.2500.20 | 2220.3 | 1055.4 | 1055.4 | 0.0 | 1041.5 | -1.3 | 1052.3 | -0.3 |
| 12.2500.35 | 1887.8 | 898.7 | 897.7 | -0.1 | 895.9 | -0.3 | 891.7 | -0.8 |
| 25.2500.5 | 1785.4 | 213.6 | 213.6 | 0.0 | 215.9 | +1.1 | 214.6 | +0.5 |
| 25.2500.20 | 2369.6 | 1137.7 | 1138.0 | +0.0 | 1164.3 | +2.3 | 1275.7 | +12.1 |
| 25.2500.35 | 2561.8 | 1268.5 | 1268.1 | -0.0 | 1300.6 | +2.5 | 1295.8 | +2.2 |
| 50.2500.5 | 1783.8 | 227.4 | 227.4 | 0.0 | 228.1 | +0.3 | 227.4 | 0.0 |
| 50.2500.20 | 2183.9 | 759.1 | 762.0 | +0.4 | 761.9 | +0.4 | 759.8 | +0.1 |
| 50.2500.35 | 2651.6 | 1164.3 | 1168.0 | +0.3 | 1177.5 | +1.1 | 1185.1 | +1.8 |
| MAMMALS | 6018.4 | 5828.2 | 5871.5 | +0.7 | 6156.3 | +5.6 | 6003.2 | +3.0 |
| PRIMATES | 4404.4 | 4218.6 | 4213.8 | -0.1 | 4629.4 | +9.7 | 4353.5 | +3.2 |
| VIROIDS | 219.9 | 211.9 | 221.5 | +4.5 | 221.1 | +4.3 | 224.5 | +6.0 |

**Table 2.**
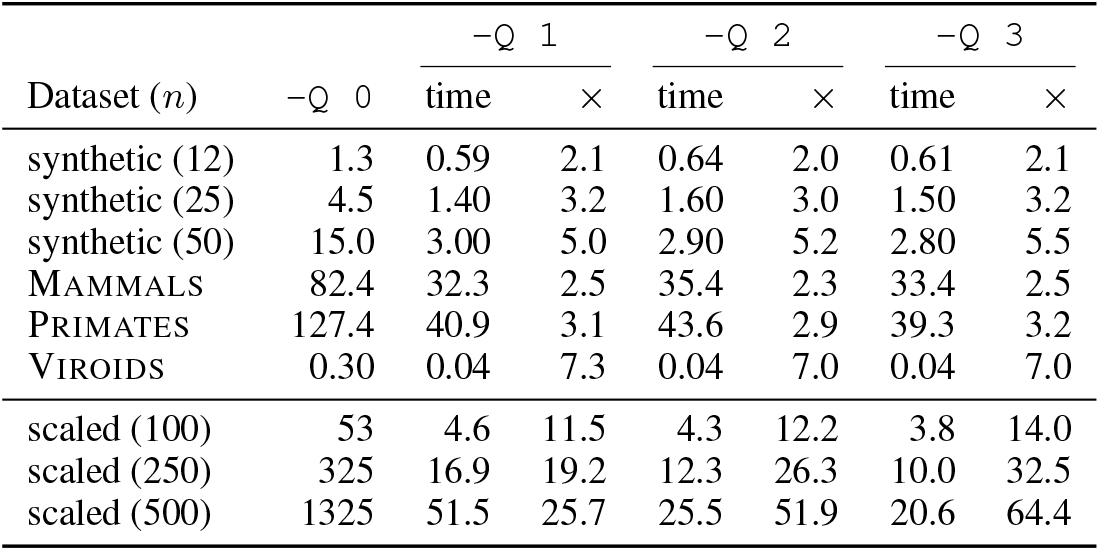
MARS wall-time (s), mean of 3 repetitions, and speedup over the reference (`-Q 0`). Synthetic rows grouped by sequence count *n*; scaled rows averaged over `med`/`high` (point values differ by up to 20% between divergence levels at the most aggressive presets, reflecting within-preset variance rather than timing noise; the per-rep run-to-run spread of every cell is below 25%, see §2.4).

| Dataset ($n$) | $-Q\ 0$ | $-Q\ 1$ | | $-Q\ 2$ | | $-Q\ 3$ | |
| --- | --- | --- | --- | --- | --- | --- | --- |
| | | time | × | time | × | time | × |
| synthetic (12) | 1.3 | 0.59 | 2.1 | 0.64 | 2.0 | 0.61 | 2.1 |
| synthetic (25) | 4.5 | 1.40 | 3.2 | 1.60 | 3.0 | 1.50 | 3.2 |
| synthetic (50) | 15.0 | 3.00 | 5.0 | 2.90 | 5.2 | 2.80 | 5.5 |
| MAMMALS | 82.4 | 32.3 | 2.5 | 35.4 | 2.3 | 33.4 | 2.5 |
| PRIMATES | 127.4 | 40.9 | 3.1 | 43.6 | 2.9 | 39.3 | 3.2 |
| VIROIDS | 0.30 | 0.04 | 7.3 | 0.04 | 7.0 | 0.04 | 7.0 |
| scaled (100) | 53 | 4.6 | 11.5 | 4.3 | 12.2 | 3.8 | 14.0 |
| scaled (250) | 325 | 16.9 | 19.2 | 12.3 | 26.3 | 10.0 | 32.5 |
| scaled (500) | 1325 | 51.5 | 25.7 | 25.5 | 51.9 | 20.6 | 64.4 |

#### Quality

Table 1 reports AVPD for the 12 original datasets. The library-scale preset stays within 1% of the original MARS on the two large real mitochondrial genomes (Mammals +0.7%, Primates -0.1%) and on every synthetic set but one, where it is 2.8% *lower* (i.e. better) than the reference; the only degradation is the short, highly divergent Viroids set (+4.5%). The aggressive presets are within roughly 1% of the original MARS on synthetic data (median |Δ| of 1.1%/0.8%, with one +12% outlier) but drift 3–10% on the large, highly divergent real genomes (Mammals, Primates) and also on the short, divergent Viroids set, the known short/high-divergence corner. Crucially, *all* presets keep AVPD far below the no-MARS baseline, so the rotation refinement still helps.

#### Runtime and speedup

Table 2 shows MARS wall-time (mean of 3 reps) and the speedup over the reference. At the paper’s scale (up to 50 sequences) the speedup is a modest constant factor of 2–7× (the short Viroids set aside, which already benefits more). This is expected: at small sequence counts the removed quadratic and cubic stages are not yet dominant, so the gains reflect only the constant-factor wins (the dropped refinement and the faster tree); the scaling improvement only becomes visible at large counts. The scaled rows of Table 2 (discussed in §3.2) show this clearly, with speedup growing to well over an order of magnitude at 500 sequences.

### 3.2 BENCHMARK ON SYNTHETIC SCALED DATASETS

The scaled synthetic sets serve a specific purpose: to expose the large-library regime, where the removed quadratic and cubic stages dominate and the scaling gains become visible, which the original benchmark (capped at 50 sequences) cannot show. The speedup over the reference grows with the sequence count to well over an order of magnitude at 500 sequences (Table 2, scaled rows; Figure 1), reflecting the removal of those stages. Figure 2 confirms the mechanism: at 500 sequences the pairwise and refinement phases collapse, leaving the progressive alignment as the floor that all presets share.

**Figure 1.**
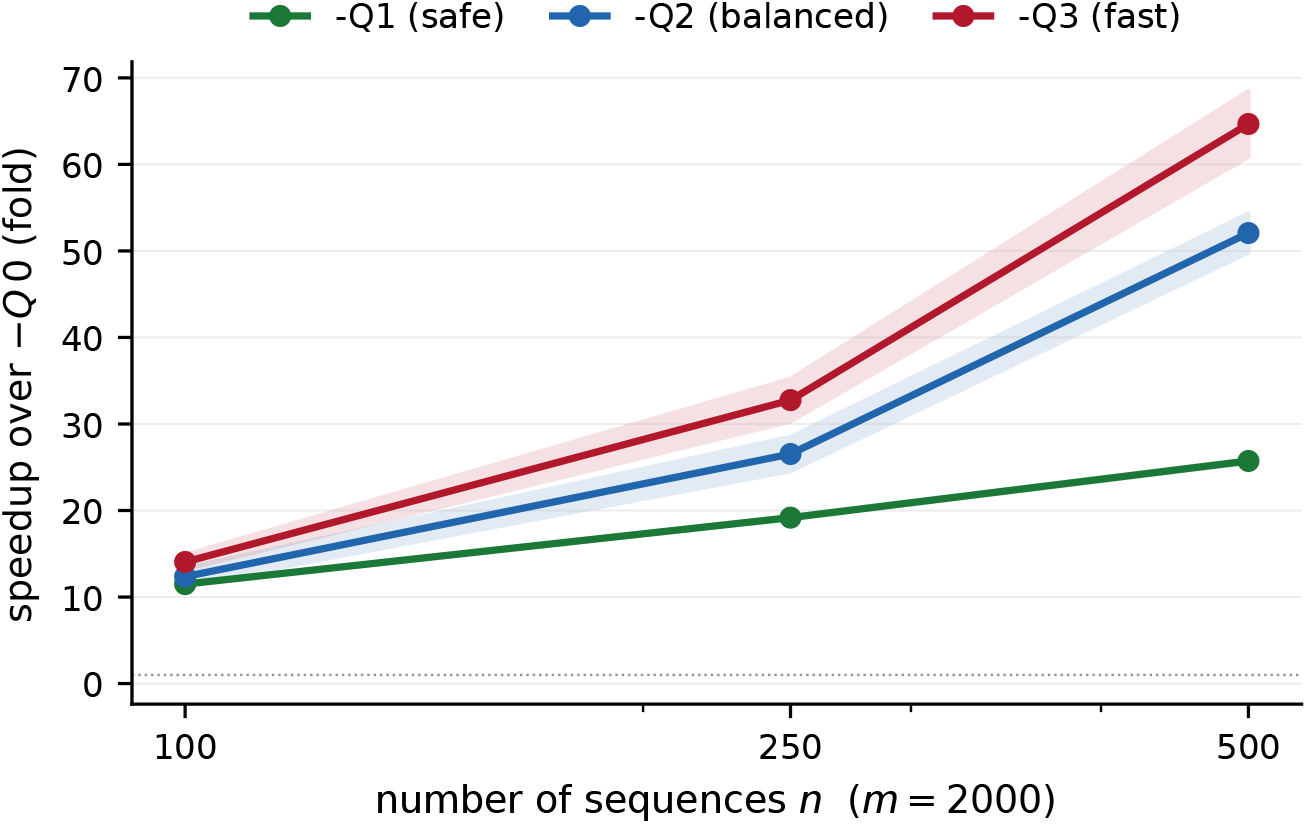
Speedup over the reference (`-Q 0`) on the scaled sets (sequence length 2000), mean of `med`/`high` with the range shaded. Speedup grows with the sequence count as the quadratic and cubic stages are removed.

**Figure 2.**
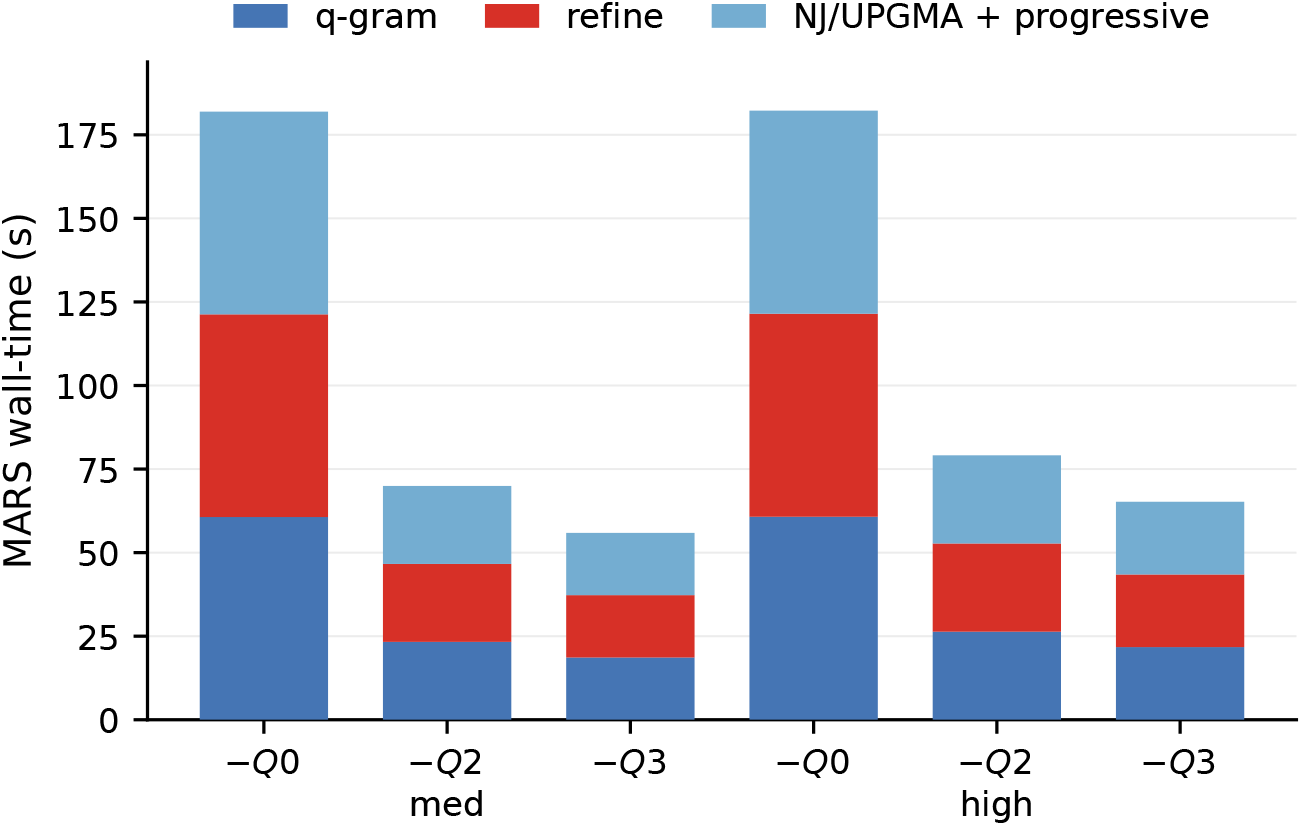
Per-phase MARS wall-time at 500 sequences (length 2000). Under the optimised presets the pairwise and refinement phases collapse; the remaining cost is the progressive alignment that all presets share.

Table 3 and Figure 3 report quality at scale. At moderate divergence the library-scale preset and `-Q 3` match the reference closely (AVPD within *∽* 1.5%, circvar *≈* 10^−4^, rotation agreement *≈* 1.000); the most aggressive preset (`-Q 2`) drifts more on the largest set (`n500.med`, +5.3%, circvar 0.0053). The only genuine soft spot is the largest, most divergent set, *n*=500 `high`, where the library-scale preset’s guide tree and rotation vote lose the frame for a minority of sequences (circvar 0.014, AVPD +5.8%); the aggressive presets stay closer to the reference there.

**Table 3.** Scaled sets (length 2000): AVPD / circvar / rotation agreement (rotagr) vs the reference (-Q 0). circvar 0 = perfect frame (*∽* 0.95 = random); rotagr 1.0 = matches the reference up to the global frame.

| dataset | $n$ | AVPD / circvar / rotagr | | | |
| --- | --- | --- | --- | --- | --- |
| | | $-Q\ 0$ | $-Q\ 1$ | $-Q\ 2$ | $-Q\ 3$ |
| n100.med | 100 | 830.4/.0001/1.000 | 841.8/.0001/1.000 | 858.9/.0000/1.000 | 850.9/.0003/1.000 |
| n100.high | 100 | 1105.2/.0002/1.000 | 1099.3/.0002/1.000 | 1120.9/.0006/.999 | 1121.8/.0001/1.000 |
| n250.med | 250 | 902.9/.0002/1.000 | 896.3/.0001/1.000 | 915.1/.0007/1.000 | 905.7/.0001/1.000 |
| n250.high | 250 | 1217.7/.0001/1.000 | 1212.6/.0004/1.000 | 1239.4/.0015/.999 | 1249.0/.0022/.998 |
| n500.med | 500 | 998.6/.0002/1.000 | 994.1/.0001/1.000 | 1051.4/.0053/.994 | 1003.1/.0004/1.000 |
| n500.high | 500 | 1257.1/.0001/1.000 | 1330.1/.0140/.986 | 1300.8/.0060/.994 | 1278.1/.0017/.999 |

**Figure 3.**
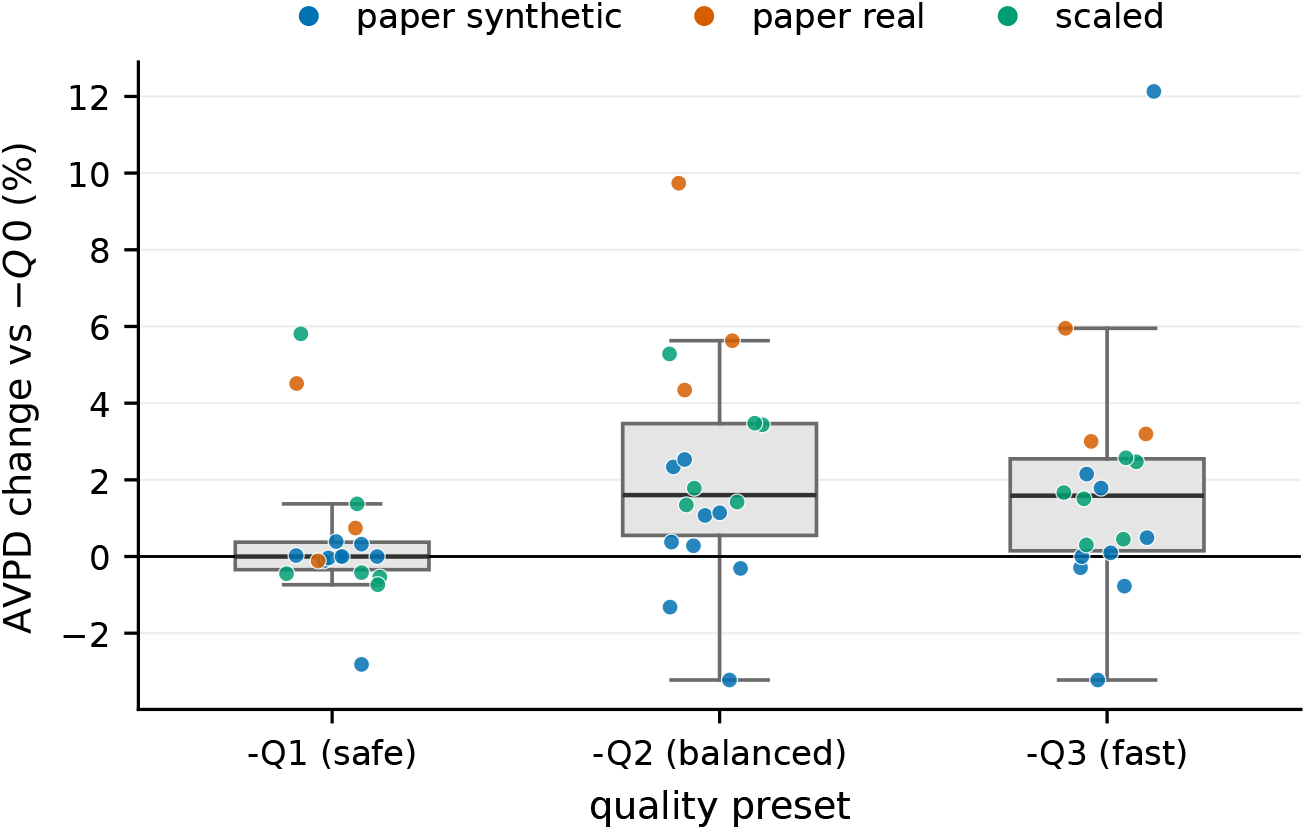
AVPD change vs the reference (`-Q 0`) across all datasets. The library-scale preset clusters at zero on synthetic and scaled-moderate data, with a small tail on the short, highly divergent Viroids and `n500.high` sets; the aggressive presets show the same pattern more strongly.

### 3.3 H2T CYCLIC PEPTIDE LIBRARY: MOTIF RECsOVERY UNDER ROTATION

To test the reference-based rotation-recovery scheme (§2.3B) on realistic inputs rather than synthetic controls, we built a library of 1,000 H2T cyclized peptides of length 18 targeting MDM2 produced by an internal peptide-design engine. Each sequence was then cyclically rotated by an independent random offset, destroying any shared frame and giving MARS a genuine recovery task. As a baseline we aligned the rotated stack directly with ClustalΩ (which cannot re-frame cyclic sequences): no recognisable motif is recoverable, reflected both in a high average pairwise distance and in an essentially flat information-content profile (Figure 4, top). Applying MARS restores a common frame and the designed positional conservation reappears as tall, column-aligned stacks (Figure 4, Q0/Q1). Crucially, the library-scale preset recovers the same motif as the original reference while running about 1.4 s versus about 942 s (mean *±* SEM over 3 replicates), an end-to-end speedup of over 650× . On the column-comparable ClustalΩ-AVPD it scores 14.39 versus the reference’s 13.67, retaining approximately 92% of the reference’s improvement over the unaligned baseline (22.62). On the stricter same-length Hamming AVPD (the mean pairwise Hamming distance on the ungapped, equal-length rotated stack, computed without any alignment step) the library-scale preset retains about 66% of the reference’s gain (13.78 vs 12.43, baseline 16.46). Table 4 reports the per-preset quality and runtime, and Figure 4 the information-content logos. This makes motif discovery by MCSA practical at generative-model library scale (1,000 sequences in about 1.4 s).

**Table 4.** H2T cyclic peptide library (1,000 sequences, length 18, target MDM2; randomly rotated input). ClustalΩ-AVPD is computed on the final MSA; Hamming-AVPD is the mean pairwise Hamming distance on the ungapped same-length stack (no alignment). Runtime is mean *±* SEM over 3 reps on an Intel Xeon 6.

| | MSA len | ClustalΩ-AVPD | Hamming-AVPD | runtime (s) |
| --- | --- | --- | --- | --- |
| baseline (no MARS) | 67 | 22.62 | 16.46 | — |
| $-Q\ 0$ (reference) | 37 | 13.67 | 12.43 | 942.09 ± 2.4 |
| $-Q\ 1$ (recommended) | 45 | 14.39 | 13.78 | 1.365 ± 0.003 |

**Figure 4.**
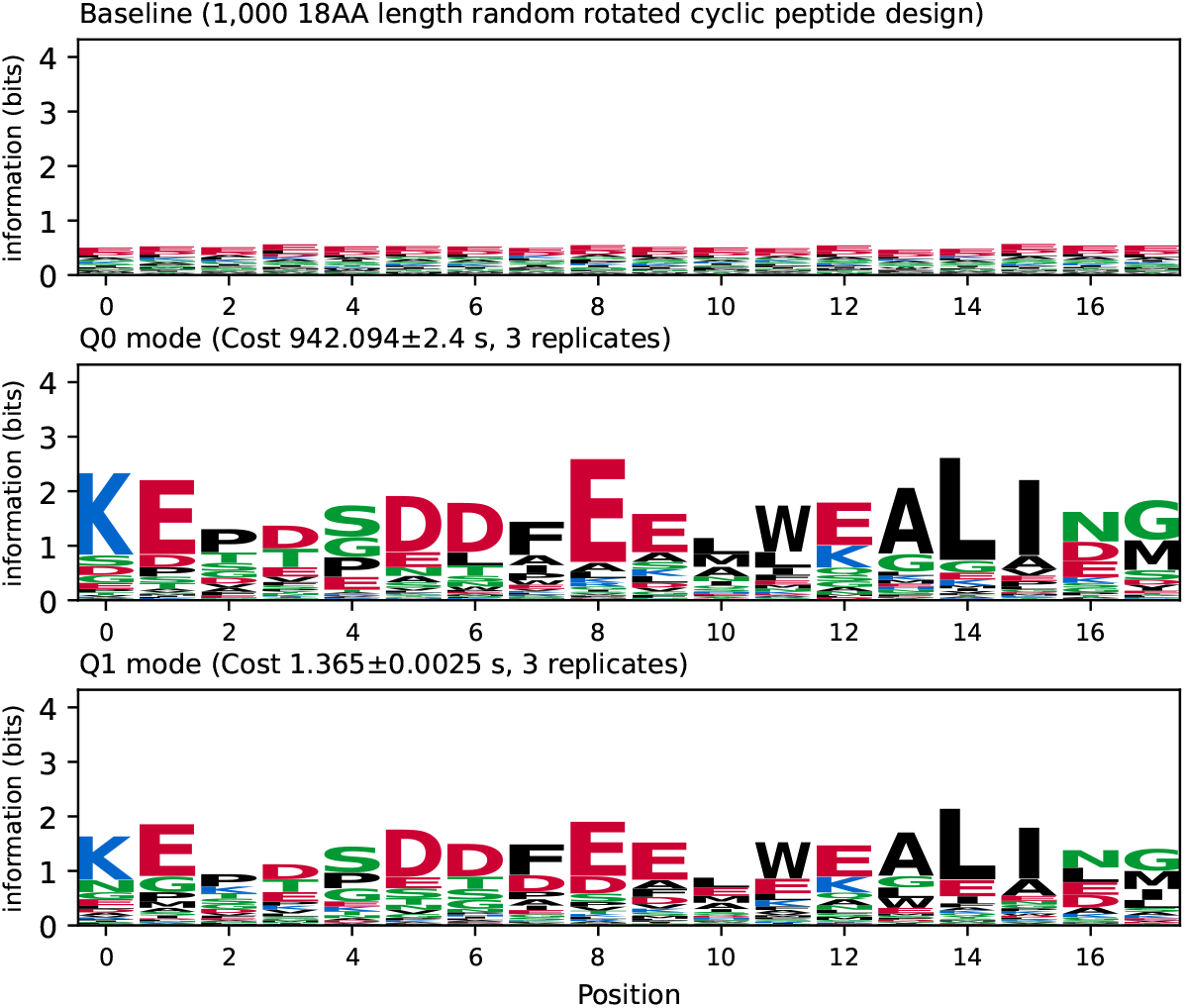
Information-content sequence logos for the cyclic peptide library (1,000 sequences) under random per-sequence cyclic rotation. Top: the baseline, a ClustalΩ alignment of the rotated stack with no cyclic re-framing; the profile is flat, showing no recoverable motif. Middle/Bottom: MARS `-Q 0` (original reference) and `-Q 1` (library-scale preset), with `-Q 1` brought into `-Q 0`’s frame by a single global cyclic shift minimising the *L*_2_ distance between position-frequency matrices so the two rows are column-comparable. Letter height is position information in bits; colours follow the WebLogo chemistry scheme. MARS recovers the designed motif, and `-Q 1` reproduces the `-Q 0` profile at over 650× lower runtime (mean *±*SEM, 3 replicates, in titles). CPU used for this test: Intel Xeon 6.

### 3.4 Multi-core and SIMD SCALING

The parallelisation (optimisation D) is orthogonal to the quality presets and applies to the original algorithm equally. We measure strong scaling (fixed input, increasing thread count) with the same 3-repetition protocol; run-to-run spread was under 9% for every cell (under 2% for the single-thread run). Table 5 and Figure 5 report the results.

**Figure 5.**
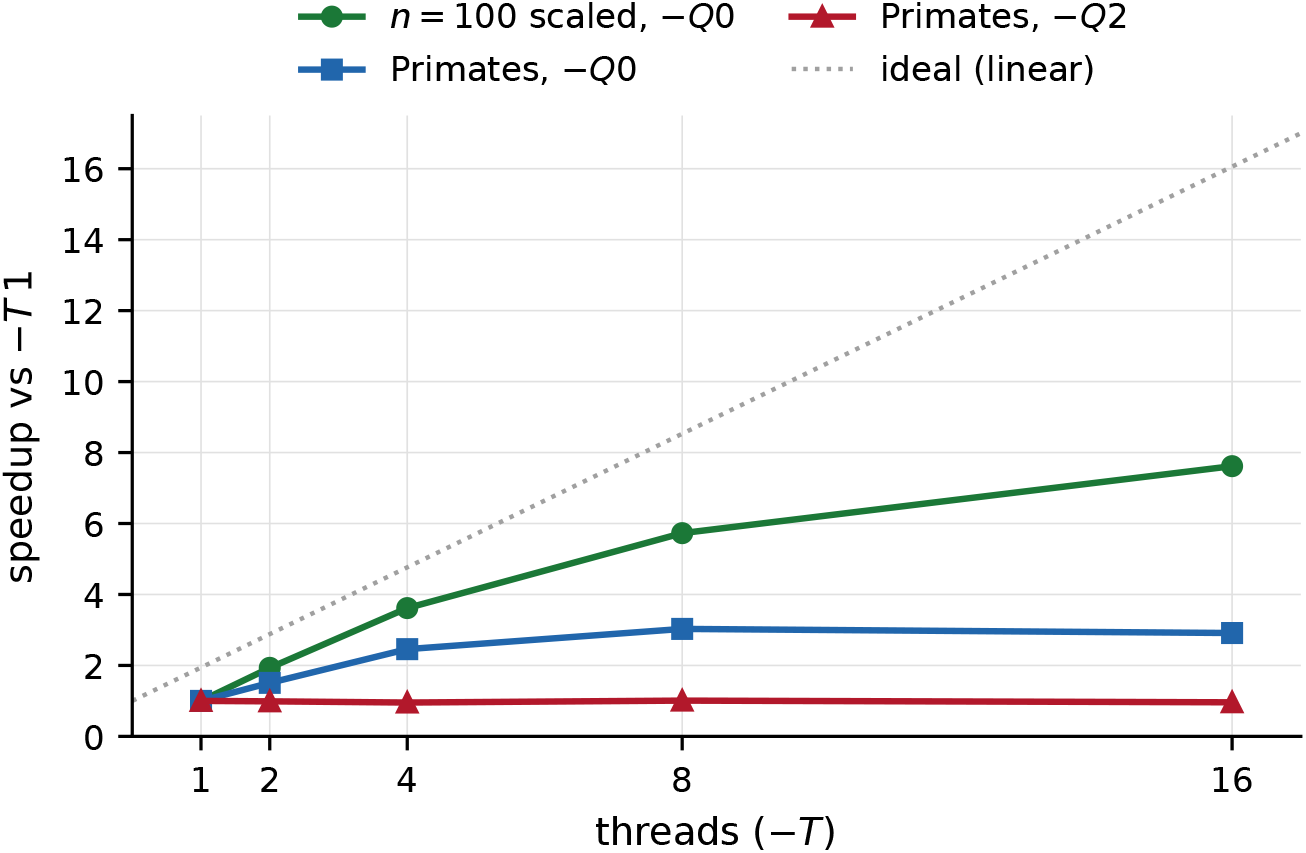
Strong scaling (speedup over the single-thread run, `-T 1`) on a multi-core node. The pairwise-comparison-bound workload (100 scaled sequences under the reference, where the original quadratic refinement dominates) scales near-linearly through 8 threads; the long-sequence Primates genome under the reference scales sub-linearly as the progressive tail comes to dominate; and the progressive-alignment-bound case (Primates under an aggressive preset, which removes the refinement stage) is limited by data dependencies and barely scales.

**Table 5.** Strong scaling: MARS wall-time (s, mean of 3 reps) and speedup over the single-thread run (-T 1), on a multi-core node. Primates = 16 real mtDNA (*∽*16 kbp); n100.med = 100 scaled sequences (length 2000).

| Dataset | $-T 1$ | | $-T 2$ | | $-T 4$ | | $-T 8$ | | $-T 16$ | |
| --- | --- | --- | --- | --- | --- | --- | --- | --- | --- | --- |
| | s | × | s | × | s | × | s | × | s | × |
| PRIMATES, $-Q 0$ | 123.2 | 1.0 | 81.7 | 1.5 | 50.1 | 2.5 | 40.7 | 3.0 | 42.3 | 2.9 |
| $n100.med$, $-Q 0$ | 52.4 | 1.0 | 27.1 | 1.9 | 14.5 | 3.6 | 9.1 | 5.7 | 6.9 | 7.6 |
| PRIMATES, $-Q 2$ | 40.0 | 1.0 | 40.5 | 1.0 | 41.9 | 1.0 | 39.7 | 1.0 | 41.6 | 1.0 |

Two regimes are visible, and both are as the design predicts. When the quadratic pairwise stage dominates (the original refinement under the reference, which accounts for most of the single-thread time at 100 short sequences), parallelism scales near-linearly through 8 threads (1.9 × at 2 threads, 3.6× at 4, 5.7× at 8, i.e. 71–95% of ideal), and continues to improve at 16 threads (7.6× ) where the data-dependent progressive tail begins to bound the speedup. On the real Primates genome (16 sequences of about 16 kbp) under the reference the pairwise refinement again dominates the single-thread time and parallelism scales sub-linearly through 8 threads (3.0×), after which the pro-gressive alignment becomes the bottleneck. The third row makes the orthogonality precise: the aggressive preset *removes* the easily-parallelised refinement, leaving the progressive alignment, whose data dependencies limit parallelism and which therefore does *not* scale with threads (the Primates aggressive-preset row is essentially flat). For long-sequence inputs the aggressive presets and the multi-core backend are therefore alternative routes to a similar wall-time; for many-sequence inputs they compose, since the reference’s large pairwise stage is exactly what parallelism excels at. The parallel output is bit-identical to single-thread on all datasets (0 differing bytes), so increasing the thread count incurs no quality cost.

## 4 Discussion

### Advantages and when to use which preset

The library-scale preset, which drops the redundant refinement and uses the faster guide tree, preserves quality to within 1% on the large real mitochondrial genomes (Mammals, Primates) and within 3% on the synthetic sets (the one exception is a 2.8% *improvement* over the reference), with the only genuine soft spot the short, highly divergent Viroids set (+4.5%). It removes the quadratic refinement and the cubic guide tree, and is therefore the recommended default for cyclic peptide libraries. The aggressive presets, which additionally estimate pairwise relations from a few references, are appropriate when maximum speed is needed *and* the input is not both large and highly divergent; in that hard regime the library-scale preset should be preferred. The bit-identical multi-core and SIMD backend further reduces the remaining wall-clock through near-linear thread scaling on the pairwise-comparison stage, and composes with the algorithmic presets on many-sequence inputs.

### Limitations

We use a single downstream MSA program (ClustalΩ); absolute AVPD values therefore differ from the 2017 paper’s ClustalΩ version, but the AVPD *formula* is exact-validated against the published MUSCLE alignments and all presets use the same program, so the comparison is internally controlled. We do not report phylogenetic Robinson–Foulds distances or consistency scores (out of scope); AVPD is the sole quality metric, consistent with the original paper’s time–accuracy table. Every timing was averaged over three repetitions with strict run-to-run spread thresholds (none flagged), so the reported speedups reflect algorithmic scaling rather than hardware variability. Our claim that the progressive alignment is close to optimal (Supplementary Material, Table S2) is a conditional lower bound, not an unconditional one, and does not preclude an approximate aligner from doing better. The aggressive presets measurably degrade on highly divergent real mtDNA and at 500 sequences of high divergence; we have characterised rather than hidden this trade-off.

### Future directions

Several directions follow naturally from the current limitations. The remaining bottleneck is the progressive alignment itself, which is already close to a theoretical limit for exact methods (Supplementary Material); an *approximate* aligner could in principle go faster, though no current MCSA tool does so. The guide-tree stage can likely be made faster still via landmark clustering (Blackshields et al., 2010). On the evaluation side, tuned presets for short peptides (§3.3) remain to be explored. Finally, the hardware route (multi-core and SIMD, and beyond it GPU) reduces the constant cost without changing the scaling.

## 5 Conclusions

The quadratic pairwise stage and the cubic guide-tree stage of MARS are not intrinsic to the MCSA problem: they can be reduced to near-linear cost in the sequence count without changing the output meaningfully. The result is one to two orders of magnitude of end-to-end speedup with quality preserved for the library-scale preset, making multiple circular sequence alignment practical at the library sizes (hundreds to thousands of sequences) relevant to cyclic peptide drug discovery for the first time. Combined with a reference-based rotation-recovery scheme that, to our knowledge, is the first such acceleration specific to circular sequences, and a bit-identical multi-core and SIMD backend, this turns MCSA from a tool that takes minutes on a dozen sequences into one that handles hundreds in seconds, suitable as a motif-discovery front end for cyclic peptide generation pipelines. The optimisations ship behind a quality preset in the MARS build, and the full evaluation harness (dataset acquisition, runner, metric calibration, figure generation) is released alongside the paper.

## Author contribution

F.H., P.P. and K.H. secured funding for the project. Y.Y. conceptualised the project and implemented proposed optimisations with input from Z.L. and K.H. Z.L. performed the benchmark on internal cyclic peptide design libraries.

## Acknowledgements

This work was supported by Pharmaron’s internal R&D grant (Bio-PHAR-YFG-2603002). We thank our colleagues in the Generative Biology Group, In Vitro Biology, Pharmaron, for their input and advice.

## Generative ai statement

Generative AI (GLM 5.2) was used to facilitate coding and literature search for this project. AI-generated code and AI-identified literature were manually verified by the author. Extensive benchmarking and validation further demonstrate reliability of the main contributions.

## Footnotes

1 Run-to-run spread never exceeded 25% on any of the 72 timed cells; the longest cell (n=500 under the reference preset, ∽22 min) varied by under 0.1%.

